# Evolution of an enzyme from a solute-binding protein

**DOI:** 10.1101/157495

**Authors:** Ben E. Clifton, Joe A. Kaczmarski, Paul D. Carr, Monica L. Gerth, Nobuhiko Tokuriki, Colin J. Jackson

## Abstract

Much of the functional diversity observed in modern enzyme superfamilies originates from molecular tinkering with existing enzymes^1^. New enzymes frequently evolve from enzymes with latent, promiscuous activities^2^, and often inherit key features of the ancestral enzyme, retaining conserved catalytic groups and stabilizing analogous intermediates or transition states^3^. While experimental evolutionary biochemistry has yielded considerable insight into the evolution of new enzymes from existing enzymes^4^, the emergence of catalytic activity *de novo* remains poorly understood. Although certain enzymes are thought to have evolved from non-catalytic proteins^5–7^, the mechanisms underlying these complete evolutionary transitions have not been described. Here we show how the enzyme cyclohexadienyl dehydratase (CDT) evolved from a cationic amino acid-binding protein belonging to the solute-binding protein (SBP) superfamily. Analysis of the evolutionary trajectory between reconstructed ancestors and extant proteins showed that the emergence and optimization of catalytic activity involved several distinct processes. The emergence of CDT activity was potentiated by the incorporation of a desolvated general acid into the ancestral binding site, which provided an intrinsically reactive catalytic motif, and reshaping of the ancestral binding site, which facilitated enzyme-substrate complementarity. Catalytic activity was subsequently gained *via* the introduction of hydrogen-bonding networks that positioned the catalytic residue precisely and contributed to transition state stabilization. Finally, catalytic activity was enhanced by remote substitutions that refined the active site structure and reduced sampling of non-catalytic states. Our work shows that the evolutionary processes that underlie the emergence of enzymes by natural selection in the wild are mirrored by recent examples of computational design and directed evolution of enzymes in the laboratory.

## Main text

Solute-binding proteins (SBPs) comprise an abundant and adaptable superfamily of extracytoplasmic receptors that are mainly involved in solute transport and chemotaxis in association with bacterial ATP-binding cassette (ABC) importers and chemotactic receptors^8^. However, enzymes such as cyclohexadienyl dehydratase (CDT; EC 4.2.1.51, 4.2.1.91), which catalyzes the cofactor-independent Grob-type fragmentation of prephenate and L-arogenate to yield phenylpyruvate and L-phenylalanine^9^, have apparently evolved from this superfamily of non-catalytic proteins (**Fig. 1a and Supplementary Table 1**). The relationship between CDTs and SBPs was initially recognized based on sequence similarity between CDTs and polar amino acid-binding proteins (AABPs)^5^. More recently, crystal structures of CDT from *Pseudomonas aeruginosa* (PaCDT) and a putative AABP from *Wolinella succinogenes* (Ws0279, 26% sequence identity) from structural genomics projects have further supported the close evolutionary relationship between CDTs and AABPs^10,11^. The periplasmic binding protein-like (II) fold shared by PaCDT and Ws0279 consists of two α/β domains connected by two flexible hinge strands, with the ligand binding site located at the interface of the two domains (**Fig. 1b**). Ws0279 has been annotated as a lysine-binding protein based on homology, which we confirmed using differential scanning fluorimetry (DSF) (**Extended Data Fig. 1a**).

**Figure 1.**
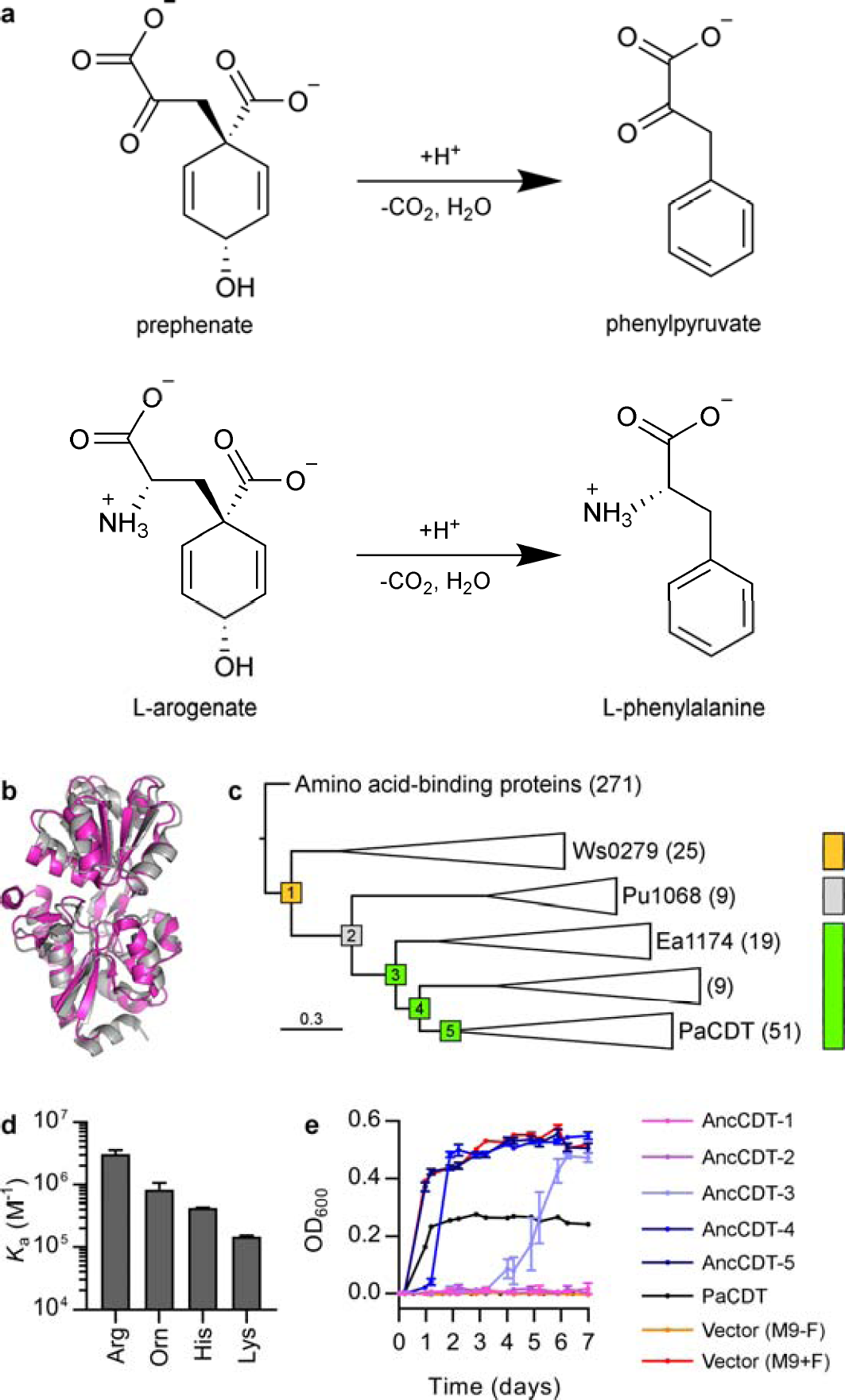
Functional evolution of CDT. **a**, Fragmentation reactions of prephenate and L-arogenate catalyzed by CDT. **b**, Structural similarity between PaCDT (grey; PDB: 3KBR) and Ws0279 (pink; PDB: 3K4U) (root-mean-square deviation 2.25 Á for backbone atoms). **c**, Condensed maximum-likelihood phylogeny of CDT homologs. The scale bar represents the mean number of substitutions per site. The five compressed clades are labeled with the corresponding number of sequences and the representative extant protein characterized in this work. The five ancestral nodes that were characterized experimentally (AncCDT-1 to AncCDT-5) are labeled and colored according to function (gold, amino acid binding; grey, binding of unknown solute; green, CDT). **d**, Affinity of AncCDT-1 for cationic amino acids, determined by ITC (Orn, L-ornithine). Results are mean ± s.d. for two (Orn, Lys) or three (Arg, His) titrations. **e**, Growth of auxotrophic *E. coli ΔpheA* cells complemented with ancestral proteins or PaCDT in selective M9-F media. Growth curves of empty vector transformants in selective M9-F media and unselective M9+F media are shown as negative and positive controls, respectively. Results are mean ± s.e.m. for four biological replicates (AncCDT-5) or five biological replicates (otherwise).

**Extended Data Figure 1.**
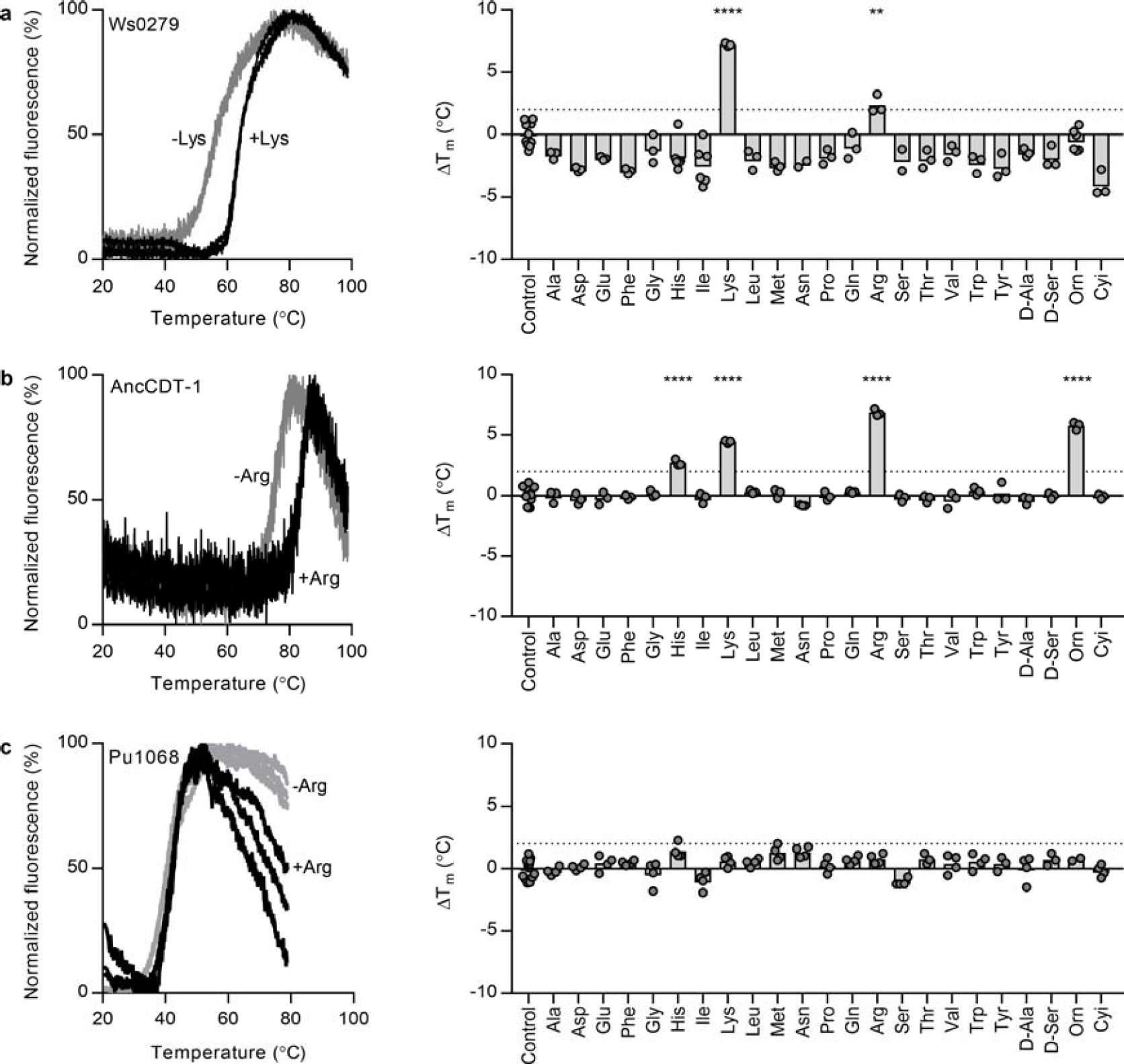
Amino acid binding profiles of Ws0279, AncCDT-1, and Pu1068. **a**, Ws0279. **b**, AncCDT-1. **c**, Pu1068. Left panels: examples of fluorescence-monitored thermal denaturation data in the absence (grey) and presence (black) of an amino acid. Three replicate curves are shown for each condition. Right panels: melting temperature (TM) of each protein in the presence of amino acids (10 mM, except for Trp, Tyr and Cyi at 1 mM), relative to a protein-only control. Columns represent the mean of the experimental replicates, shown as circles. Asterisks indicate Δ*T*_M_ > 2 °C and significantly different from the control by one-way ANOVA with Dunnett’s test for multiple comparisons (**P < 0.01, *\*\*\*\*p*< 0.0001). The Δ*T*_M_ for Ws0279 was 7.2 °C with 10 mM Lys and 6.1 °C with 1 mM Lys, comparable with Δ*T*_M_ values observed for other AABPs in the presence of their physiological ligands^63^.

To reconstruct the evolutionary history of CDT, we inferred the maximum-likelihood phylogeny of 131 homologs of Ws0279 and PaCDT, and used ancestral protein reconstruction^12^ to infer the most likely amino acid sequence for each ancestral node in the phylogeny (**Fig. 1c and Extended Data Fig. 2**). We selected five ancestral nodes, designated AncCDT-1 to AncCDT-5, for experimental characterization based on patterns of sequence conservation in the extant sequences (**Fig. 1c**). AncCDT-1 represents the last common ancestor of Ws0279 and PaCDT, while the other ancestral nodes represent intermediates in the evolution of PaCDT from AncCDT-1.

**Extended Data Figure 2.**
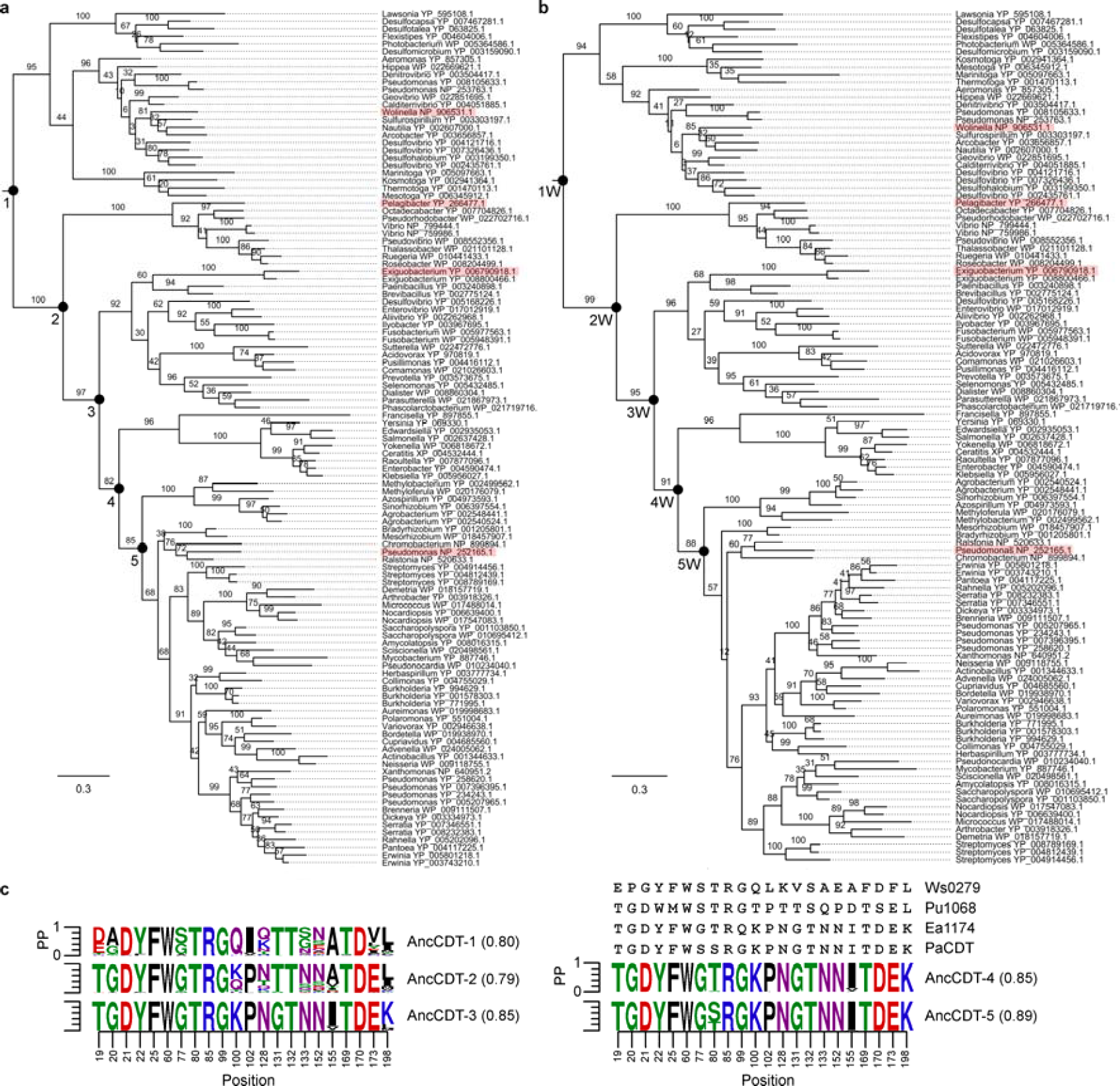
Phylogenetic analysis of CDT homologs. **a-b**, Maximum-likelihood phylogenies inferred using the LG substitution matrix (a) and the WAG substitution matrix (b). Branches are labeled with bootstrap values from 100 replicates. For each protein sequence, the NCBI accession code and the genus of the source organism are given. Experimentally characterized extant proteins are highlighted, and experimentally characterized ancestral nodes are labeled. The scale bar represents the mean number of substitutions per site. The outgroup of 271 AABP sequences is not shown. **c**, Posterior probability distributions of ancestral protein sequences at positions important for amino acid binding or CDT activity, as indicated by structural analysis or directed evolution. The sequences of Ws0279, Pu1068, Ea1174, and PaCDT at the corresponding positions are shown. The mean posterior probability of each ancestral sequence is given in parentheses.

We experimentally characterized the five ancestral proteins, using isothermal titration calorimetry (ITC) to test for amino acid binding and genetic complementation to test for enzymatic activity; in the genetic complementation assay, expression of CDT rescues the growth of *Escherichia coli* L-phenylalanine auxotrophs that lack prephenate dehydratase encoded by the gene *pheA^9^.* AncCDT-1 is an amino acid-binding protein, displaying high affinity and broad specificity for cationic amino acids, including L-arginine (*K_d_* 0.32 μM), L-ornithine (1.2 μM), L-histidine (2.3 μM) and L-lysine (6.7 μM) (**Fig. 1d and Extended Data Fig. 1b**). Neither AncCDT-2 nor any subsequent ancestral protein exhibited binding of proteinogenic amino acids. AncCDT-3, AncCDT-4, and AncCDT-5 have sufficient CDT activity to rescue growth of *E. coli ΔpheA* cells in minimal media (**Fig. 1e**). To test the phenotypic robustness of the predicted ancestral sequences to variations in the phylogenetic analysis, alternative versions of the ancestral proteins, designated AncCDT-1W to AncCDT-5W, were reconstructed using an alternative evolutionary model; genetic complementation assays using these alternative ancestral proteins gave qualitatively similar results (**Extended Data Figs 2b and 3a**). However, AncCDT-3W transformants exhibited faster growth than AncCDT-3 transformants; recombination of the two genes using staggered extension PCR followed by genetic selection showed that a single substitution (P188L) in AncCDT-3 was sufficient to recapitulate the higher growth rate associated with AncCDT-3W (**Extended Data Fig. 3b**). Spectrophotometric kinetic assays *in vitro* confirmed that AncCDT-3 and AncCDT-3(P188L), but not AncCDT-2, have prephenate dehydratase activity (**Extended Data Fig. 3d–h**).

**Figure 2.**
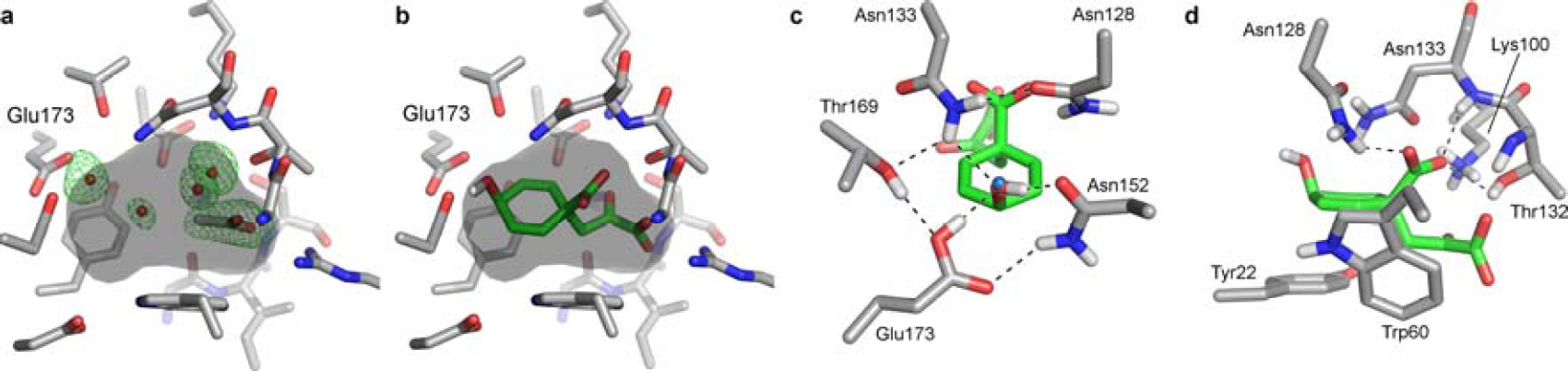
Crystal structure of PaCDT. **a**, Active site of PaCDT. The surface of the active site is shown in grey. Electron density for water and acetate molecules is shown by an omit mFM_o_ DF_c_ map contoured at +3σ. **b**, Structure of the PaCDT-prephenate complex predicted by computational docking. Docking with L-arogenate yielded a similar pose. **c**, Glu173 is poised for proton donation to the departing hydroxyl group of prephenate by hydrogen bonding interactions with neighboring residues. The position predicted to be occupied by the hydroxyl group of prephenate is occupied by a water molecule in the unliganded PaCDT structure (blue sphere). **d**, 7i-stacking interactions with Tyr22 and Trp60, and polar interactions with Lys100, Asn128, Thr132, and Asn133 could contribute to transition state stabilization.

**Extended Data Figure 3.**
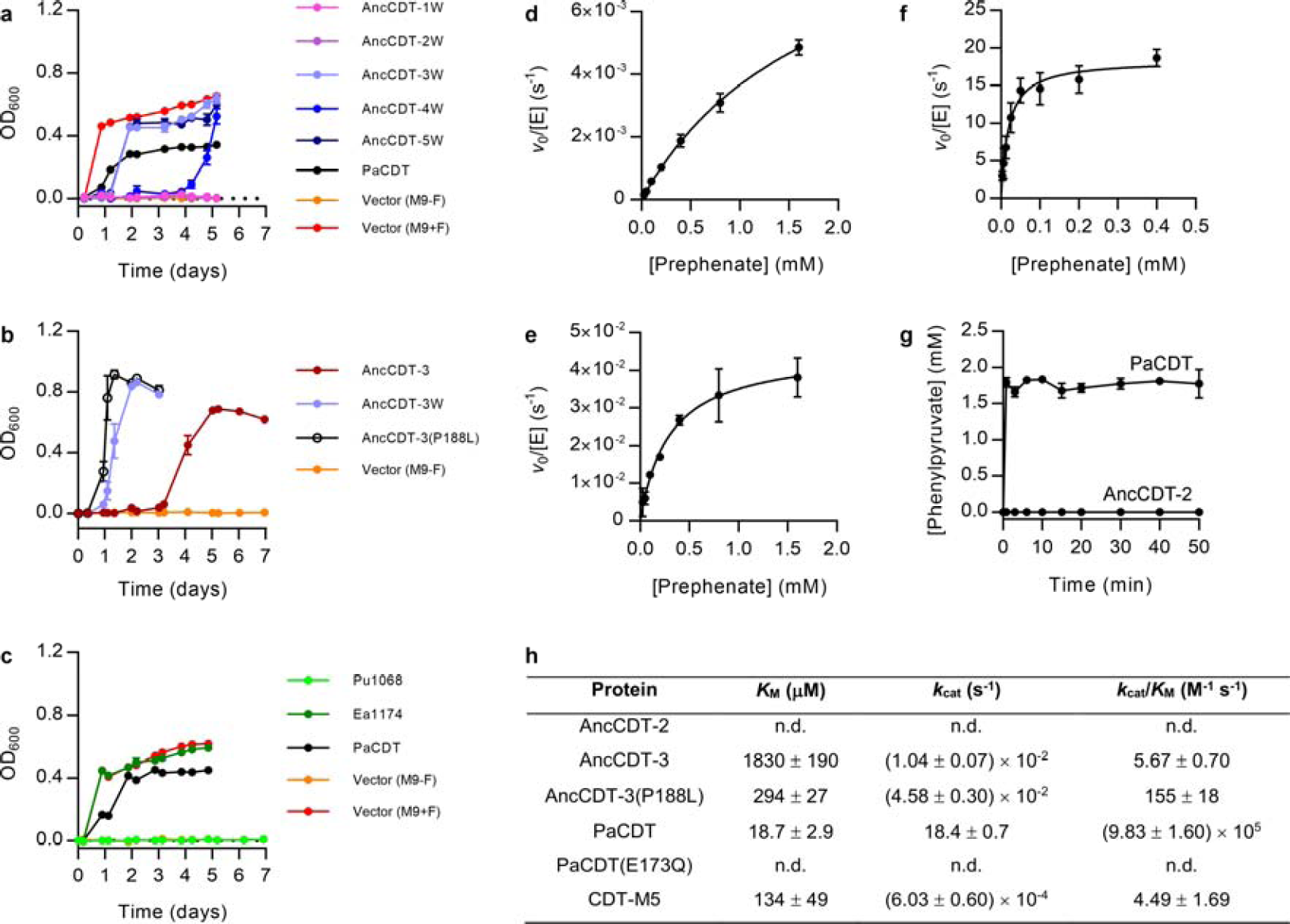
Characterization of ancestral and extant CDT variants. **a-c**, Complementation of auxotrophic *E. coli ΔpheA* cells in selective M9-F media by ancestral and extant CDT variants. Results are mean ± s.e.m. of biological replicates (a, *n* = 3; b, *n* = 5; **c**, *n* = 3). a, Alternative versions of the ancestral proteins inferred using the WAG substitution matrix (AncCDT-1 W to AncCDT-5W). b, AncCDT-3(P188L). c, Pu1068 and Ea1174. d-f, Michaelis-Menten plots for AncCDT-3 (d), AncCDT-3(P188L) (e) and PaCDT (f). Results are mean ± s.d. of technical replicates (d, *n* = 4, e, *n* = 3, f, *n* = 3). g, Conversion of 1.6 mM prephenate to phenylpyruvate by 20 M PaCDT and AncCDT-2. No activity was detected for AncCDT-2. Results are mean ± s.d., three technical replicates. h, Kinetic parameters for prephenate dehydratase activity of CDT variants characterized in this work. Errors indicate s.e. for *K*_M_ and *k*_cat_, and errors propagated from these quantities for *k*_cat_/*K*_M_. n.d., no detectable activity.

These results indicated that the ancestral amino acid-binding activity was lost between AncCDT-1 and AncCDT-2, CDT activity was gained between AncCDT-2 and AncCDT-3, and AncCDT-2 apparently had neither CDT activity nor binding affinity towards amino acids. To test whether AncCDT-2 was rendered non-functional by an error in its reconstructed sequence or had a function distinct from AncCDT-1 and AncCDT-3, we examined representatives of the previously uncharacterized evolutionary clades consisting of extant descendants of AncCDT-2 and AncCDT-3: Pu1068 from “Candidatus Pelagibacter ubique” and Ea1174 from *Exiguobacterium antarcticum* (**Fig. 1c**). Genetic complementation experiments showed that Ea1174, but not Pu1068, has CDT activity (**Extended Data Fig. 3c**), and DSF experiments showed that Pu1068 is not an amino acid-binding protein (**Extended Data Fig. 1c**). Analysis of the genomic context of *Pu1068* and several of its orthologs revealed that these genes, like the SBP gene *Ws0279,* are adjacent to genes encoding transmembrane components of ABC importers, suggesting that *Pu1068* encodes an SBP rather than an enzyme (**Supplementary Table 2**). We attempted to identify the physiological ligands of Pu1068 and AncCDT-2 *via* crystallization of Pu1068 with co-purified ligands and DSF experiments with several hundred potential metabolites from libraries and rationally selected metabolites with plausible physiological importance for oceanic bacteria such as *Ca.* P. ubique (**Extended Data Fig. 4, Supplementary Table 3**). Although the exact physiological ligands of AncCDT-2 and Pu1068 could not be identified, we found that these proteins have some affinity for a variety of carboxylates (**Extended Data Fig. 4**) and some sulfonates, such as the sulfobetaine NDSB-221, which binds Pu1068 with a *K*_d_ of 0.53 mM (**Extended Data Fig. 5**). Given that Pu1068 and its homologs are not widely distributed and are only found in bacteria that occupy a unique ecological niche (ocean), it is likely that their physiological role is highly specific for their environment and is most likely adapted for a relatively uncommon ligand. Regardless of the specific physiological ligands of AncCDT-2 and Pu1068, the functional properties of the various extant clades (Ws0279 – cationic amino acid-binding protein; Pu1068 – SBP of unknown function; Ea1174 and PaCDT – CDTs) accorded with those expected based on functional characterization of the ancestral proteins, supporting a likely evolutionary trajectory from a cationic amino acid-binding protein, to a carboxylic acid-binding protein, to CDT, an enzyme with carboxylic acid substrates (**Fig. 1c**).

**Extended Data Figure 4.**
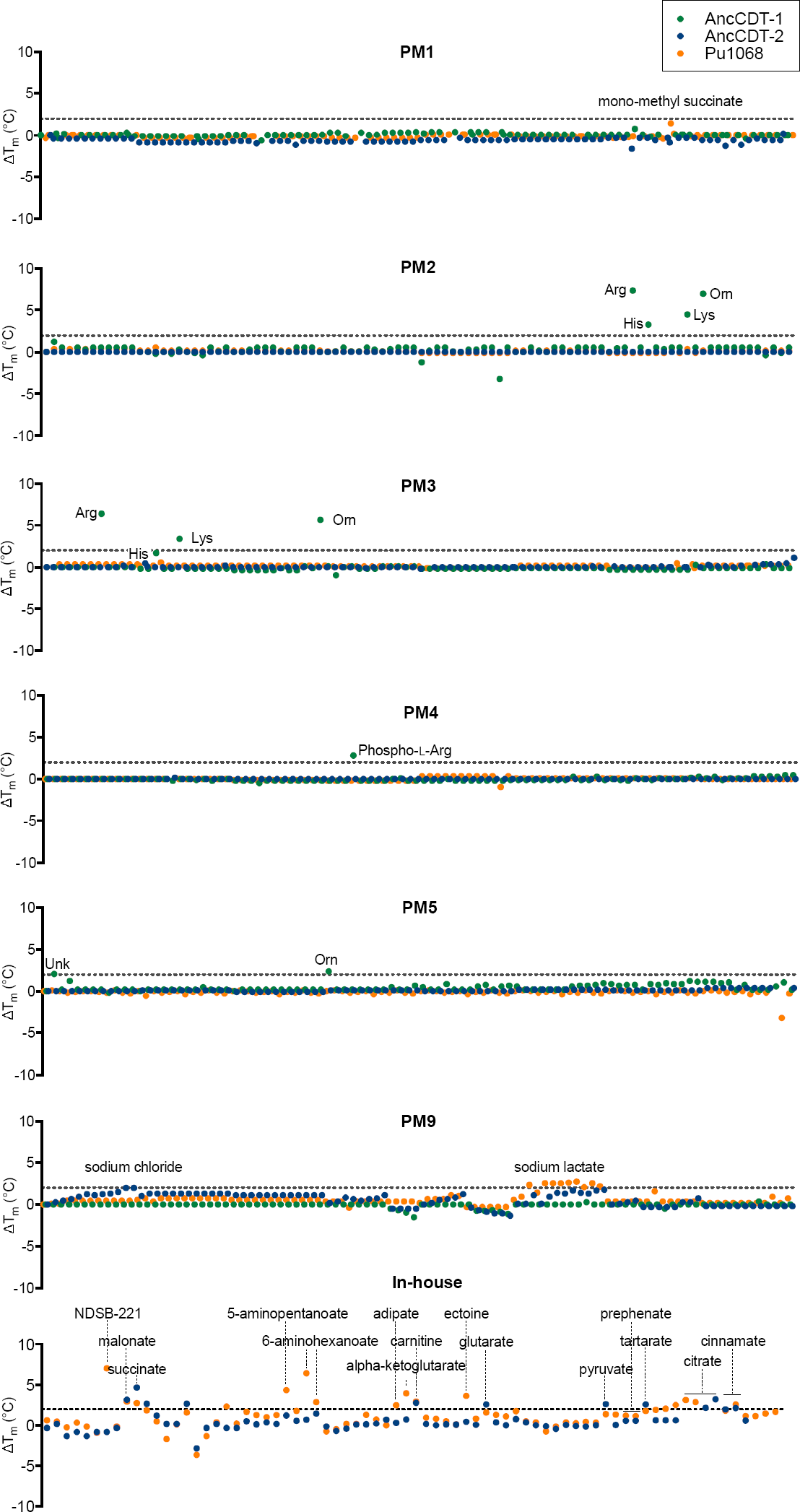
Ligand screening of Pu1068 and AncCDT-2. DSF was used to screen Pu1068, AncCDT-2, and AncCDT-1 (as a positive control) against 650 different conditions from six proprietary screens from Biolog (PM1-5 and PM9) and an in-house screen comprised of various additional compounds. Δ*T*_M_ values are given relative to a protein-only control. Compounds that produced a Δ*T*_M_ greater than 2 °C are listed. No binding of prephenate, the substrate of CDT, was observed. Details of the screen compositions and ligand concentrations are provided in **Supplementary Table 3**.

**Extended Data Figure 5.**
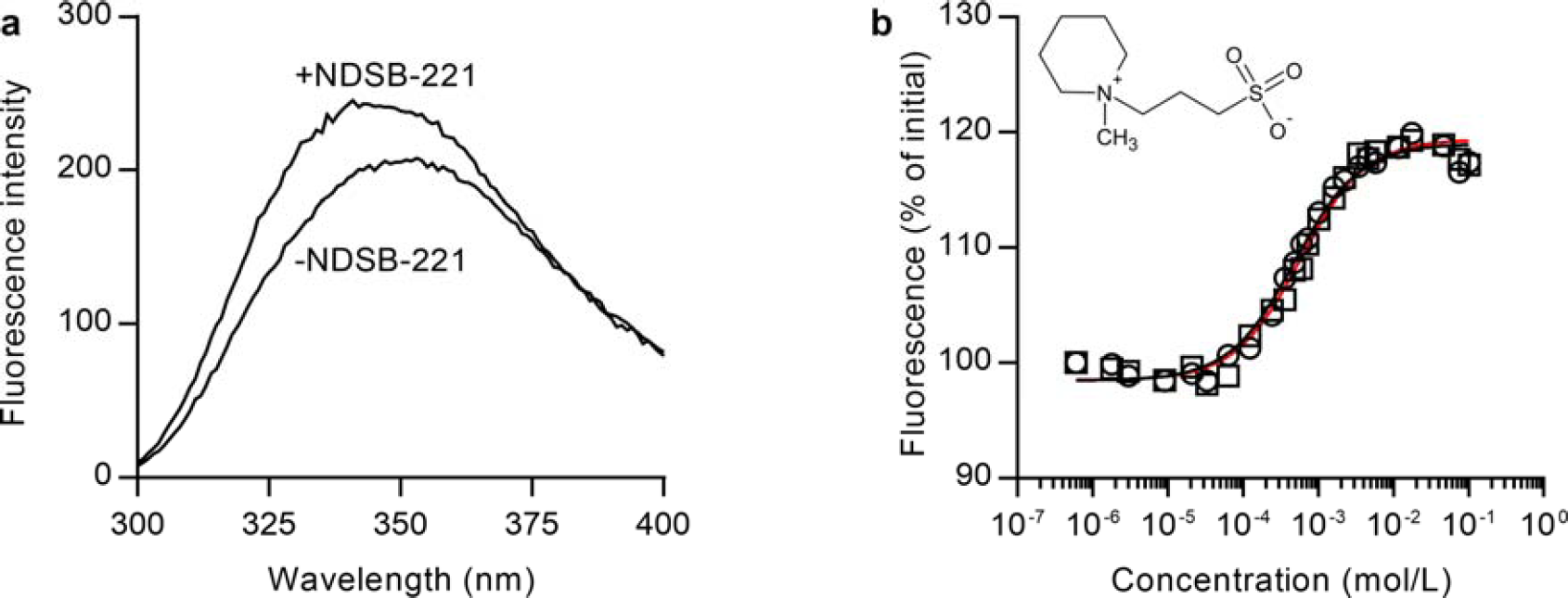
NDSB-221 is a low-affinity ligand of Pu1068. **a**, Fluorescence spectrum of Pu1068 in the presence and absence of 10 mM NDSB-221, with an excitation wavelength of 280 nm. **b**, Fluorescence titration of Pu1068 with NDSB-221; peak fluorescence is plotted against ligand concentration. Two replicate titrations are shown. Fitting the data to a Boltzmann function gives a *K_d_* of 530 M and a maximum fluorescence change of 20%. The structure of NDSB-221 is inset.

To establish the molecular basis for this functional transition, we first solved the crystal structure of unliganded PaCDT. Unlike in the crystal structure of the enzyme complexed with HEPES (**Extended Data Fig. 6**), the active site of the unliganded enzyme was fully occluded from solvent and highly complementary to its cyclohexadienol substrates (**Fig. 2a, Extended Data Fig. 6d–e**). Docking of prephenate and L-arogenate into the unliganded PaCDT structure implied a binding mode in which Glu173 is positioned adjacent to the departing hydroxyl group of the substrate, suggesting that the enzyme mechanism involves general acid catalysis by Glu173 (**Fig. 2b and Extended Data Fig. 7a**). Consistent with its proposed role as a general acid, Glu173 is partially desolvated and predicted by PROPKA to be protonated at neutral pH (p*K*_a_ 7.75), and the substitution E173Q abolishes prephenate dehydratase activity with minimal impact on secondary structure and thermostability (**Extended Data Fig. 7b–d**). The active site of PaCDT is pre-organized for protonation and elimination of the departing hydroxyl group of the substrate by an intricate hydrogen-bonding network extending from Glu173 (**Fig. 2c**). Other active site residues most likely contribute to stabilization of the departing carboxylate group and delocalized electrons in the developing π system in the transition state (**Fig. 2d**).

**Extended Data Figure 6.**
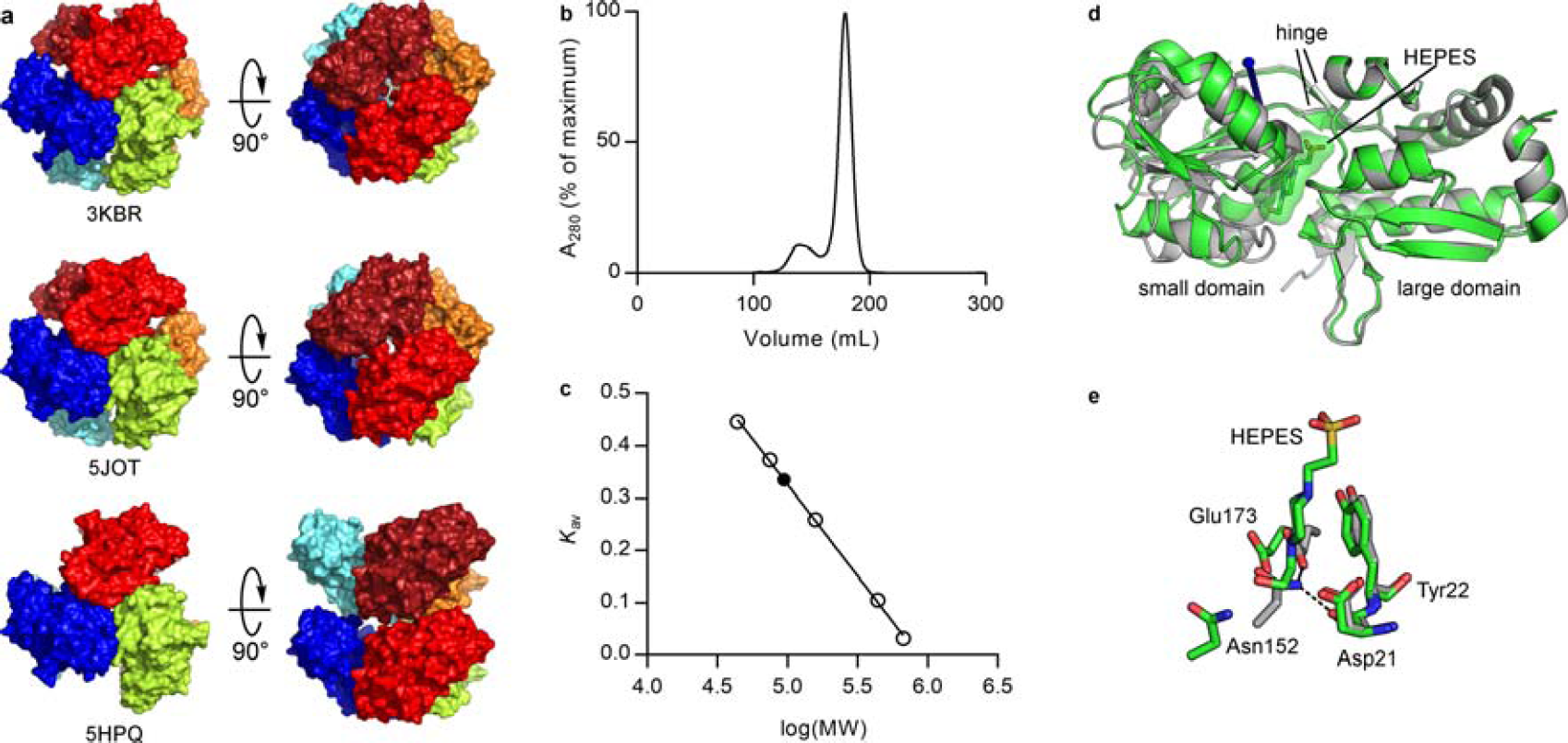
Comparison of PaCDT crystal structures. **a**, Crystallographic oligomers of PaCDT (PDB: 3KB R, 5 JOT, 5HPQ), viewed down the three-fold symmetry axis. 3KBR (HEPES-bound) and 5JOT (unliganded) show a hexameric assembly, while 5HPQ (unliganded) shows a trimeric assembly. **b**, Size-exclusion chromatogram of PaCDT. **c**, Calibration curve for analytical size-exclusion chromatography. Open circles represent molecular weight standards and the closed circle represents PaCDT. The calculated molecular weight of PaCDT is consistent with a trimeric structure (calc. 94 kDa, theor. 88 kDa for trimer). **d-e**, Conformational differences between unliganded PaCDT (PDB: 5HPQ, grey) and HEPES-bound PaCDT (PDB: 3KBR, green). d, Superimposition of the two structures using the two large domains shows a rigid-body displacement of the small domain, which corresponds to an 11° rotation about the axis indicated by the blue arrow. This conformational change accounts for occlusion of the active site in the unliganded PaCDT structure. e, HEPES disrupts the hydrogen bonding network between Asp21, Asn152, and the general acid Glu173.

**Extended Data Figure 7.**
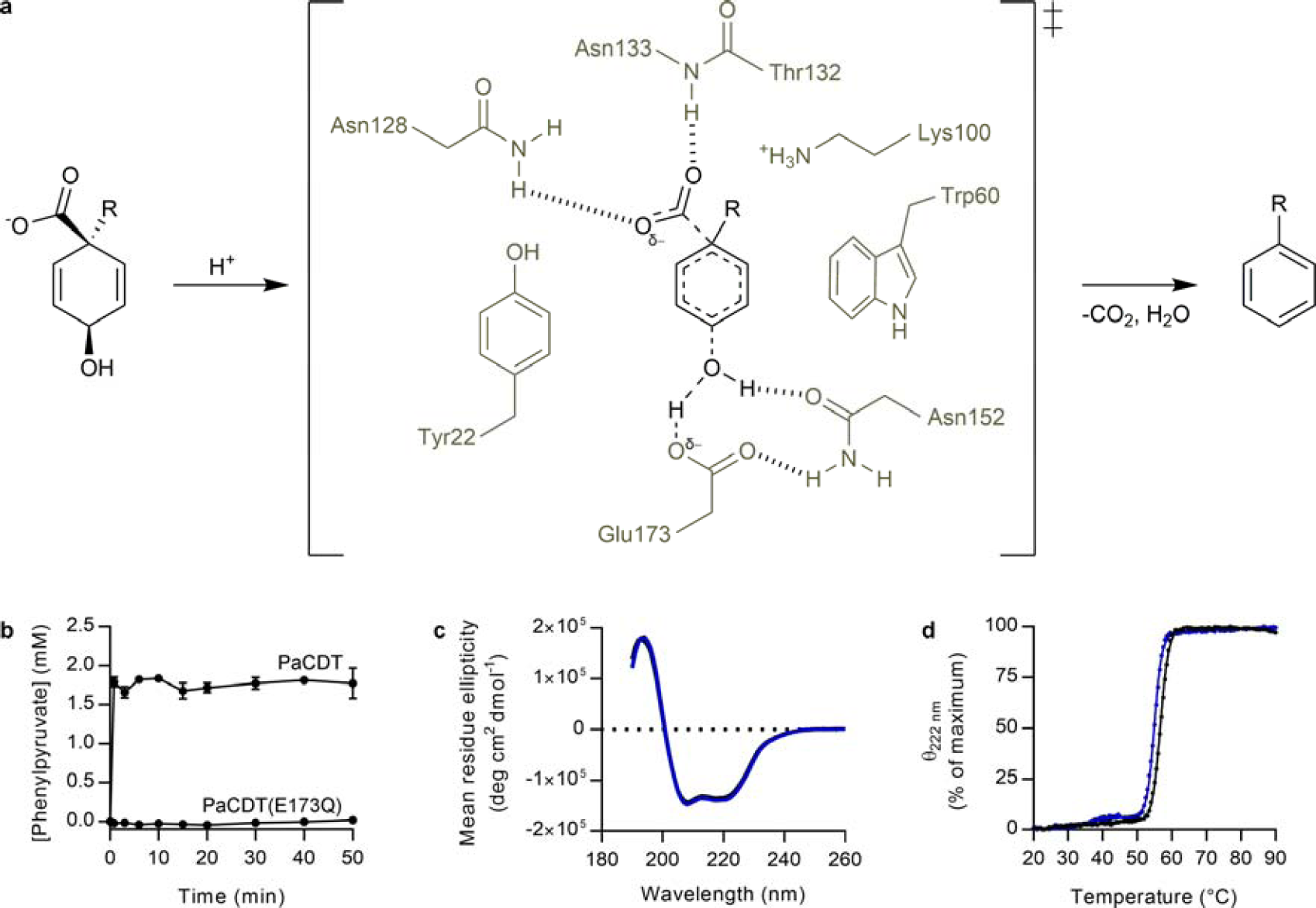
Mechanism of PaCDT. **a**, Proposed mechanism for CDT-catalyzed decarboxylative aromatization of cyclohexadienols, and basis for transition state stabilization. The general acid Glu173 donates a proton to the departing hydroxyl group of the substrate. The given mechanism shows a concerted elimination of CO2 and H2O, although stepwise elimination of H_2_O and CO_2_ *via* a divinyl carbocation intermediate is an alternative possibility. **b**, Conversion of 1.6 mM prephenate to phenylpyruvate by 20 M PaCDT and PaCDT(E173Q). The E173Q substitution abolishes prephenate dehydratase activity. Results are mean ± s.d., three technical replicates. Data for PaCDT are duplicated from Extended Data Fig. 3g; these experiments were done concurrently. **c**, Circular dichroism (CD) spectra of PaCDT (black) and PaCDT(E173Q) (blue). The E173Q substitution does not disrupt the secondary structure of PaCDT. **d**, CD-monitored thermal denaturation of PaCDT (black) and PaCDT(E173Q) (blue). The E173Q substitution has minimal impact on the *T*_M_ of PaCDT (WT, 56.6 ± 0.0 °C; E173Q, 54.9 ± 0.2 °C; mean ± s.d. for two technical replicates).

Comparison of the crystal structure of PaCDT with crystal structures of AncCDT-1, Pu1068, and AncCDT-3(P188L) revealed the contribution of historical amino acid substitutions to remodeling, functionalization, and refinement of the ancestral amino acid binding site (**Fig. 3**). Firstly, mutations that occurred between AncCDT-1 and AncCDT-2 effected two significant structural changes that potentiated the emergence of catalytic activity: the substitution V173E introduced a general acid that is positioned appropriately for general acid catalysis (**Fig. 3c**), while the substitutions D19T and A20G allowed for conformational change of Trp60, reshaping the ancestral binding site and facilitating steric complementarity between CDT and its substrates (**Fig. 3b**). These substitutions can be considered potentiating because the structural features associated with them are also observed in Pu1068 and were therefore initially adaptations towards binding a different ligand, rather than CDT activity (**Fig. 3d**). Indeed, each residue associated with these structural changes was reconstructed with high statistical confidence in the non-catalytic protein AncCDT-2 (**Extended Data Fig. 2c**). Thus, the evolution of CDT from AABPs required an intermediate adaptation to a new binding function, which introduced amino acids that potentiated the structure for subsequent evolution of catalytic function.

**Figure 3.**
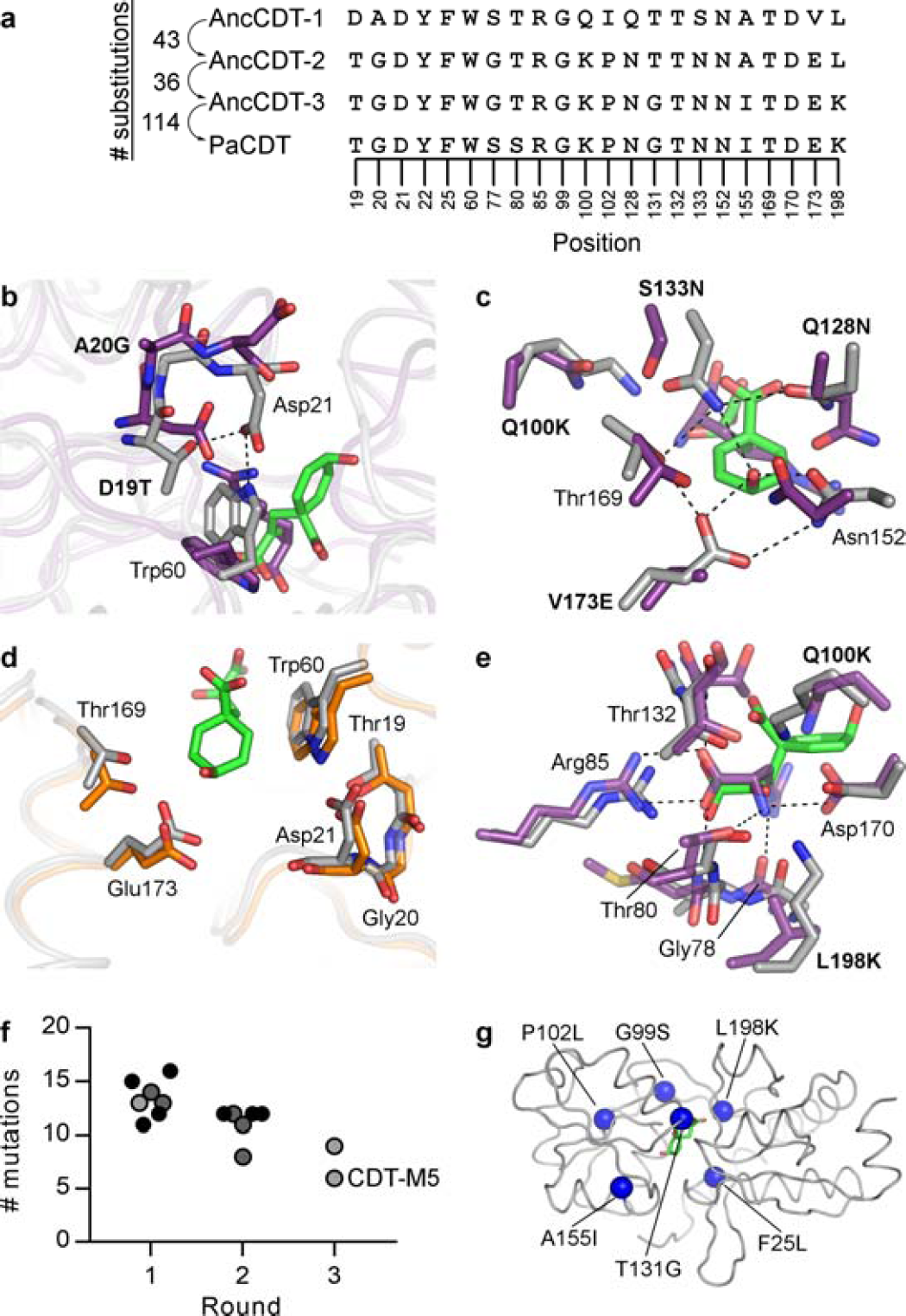
Structural and mutational basis for evolution of CDT activity. **a**, Multiple sequence alignment of ancestral proteins and PaCDT at positions important for CDT activity. The number of substitutions between each sequence in this evolutionary trajectory is shown. b-e, Comparison of AncCDT-1 (purple), Pu1068 (orange), and PaCDT (grey). Positions are labeled with the corresponding residue in AncCDT-1 and AncCDT-3, if conserved in both proteins, or with the corresponding substitution between the two proteins. **b**, The ancestral binding site was remodeled by a conformational change of Trp60. D19T introduces a hydrogen bond with Asp21, and A20G enables a conformation disfavored for non-glycine residues but necessary for the interaction between Thr19 and Asp21. **c**, Functionalization of the ancestral binding site introduced the general acid Glu173 and residues required for substrate binding and transition state stabilization. **d**, Structural similarities between Pu1068 and PaCDT. The two domains of Pu1068 were superimposed separately on the structure of PaCDT. **e**, CDT inherited the α-amino acid-binding motif from AncCDT-1, with two substitutions (Q100K, L198K) that also enable binding of the α-keto acid prephenate. **f**, Introduction of CDT activity into AncCDT-2 by directed evolution. Each point represents a unique clone, and the color gives a qualitative indication of activity (black, high activity; dark grey, moderate activity; light grey, low activity). See also Extended Data Figure 8. **g**, Positions of six substitutions sufficient to introduce CDT activity into AncCDT-2. F25L, G99S, P102L and A155I are located in the second or third shells of the active site.

Structural analysis indicates that functionalization of the ancestral binding site occurred by subsequent adaptive mutations, which fixated either between AncCDT-1 and AncCDT-2, or between AncCDT-2 and AncCDT-3. The substitutions Q100K, Q128N, and S133N introduced the hydrogen-bonding network that positions the catalytic group precisely and contributes to transition state stabilization through interactions with the departing hydroxyl and carboxylate groups of the substrate (**Fig. 3c**). Additionally, the substitutions Q100K and L198K likely contributed to dual specificity for α-amino and a-keto acid substrates (i.e., L-arogenate and prephenate) *via* electrostatic shielding of Asp170 (**Fig. 3e**). However, AncCDT-2 contains each of these four active site substitutions (except L198K, which is not itself sufficient to introduce catalytic activity) (**Fig. 3a**), implying that additional substitutions between AncCDT-2 and AncCDT-3 were required for the emergence of CDT activity. To identify these substitutions, we performed site-directed mutagenesis and three rounds of directed evolution, resulting in the isolation of an AncCDT-2 variant with only six substitutions (CDT-M5) that allowed slow growth of *E. coli* L-phenylalanine auxotrophs and exhibited prephenate dehydratase activity *in vitro* (**Fig. 3f, Extended Data Figs 3h and 8**). Although three of these substitutions (T131G, A155I, L198K) are present in AncCDT-3, the other three substitutions (F25L, G99S, P102L) represent an alternative evolutionary trajectory towards higher catalytic activity. While the T131G substitution removes a steric clash between the enzyme and the departing carboxylate group of the substrate and the L198K substitution assists binding of the ketone group, the other four substitutions are located in the second or third shells of the active site and must have indirect effects on catalysis (**Fig. 3g**). The introduction of additional mutations in various combinations supported faster growth of L-phenylalanine auxotrophs (**Fig. 3f and Extended Data Fig. 8d**). These results show that there are multiple mutational pathways to higher catalytic activity *via* remote substitutions following the introduction of key active site residues, and that the evolutionary trajectory towards high catalytic activity is not strongly deterministic in this case.

**Extended Data Figure 8.**
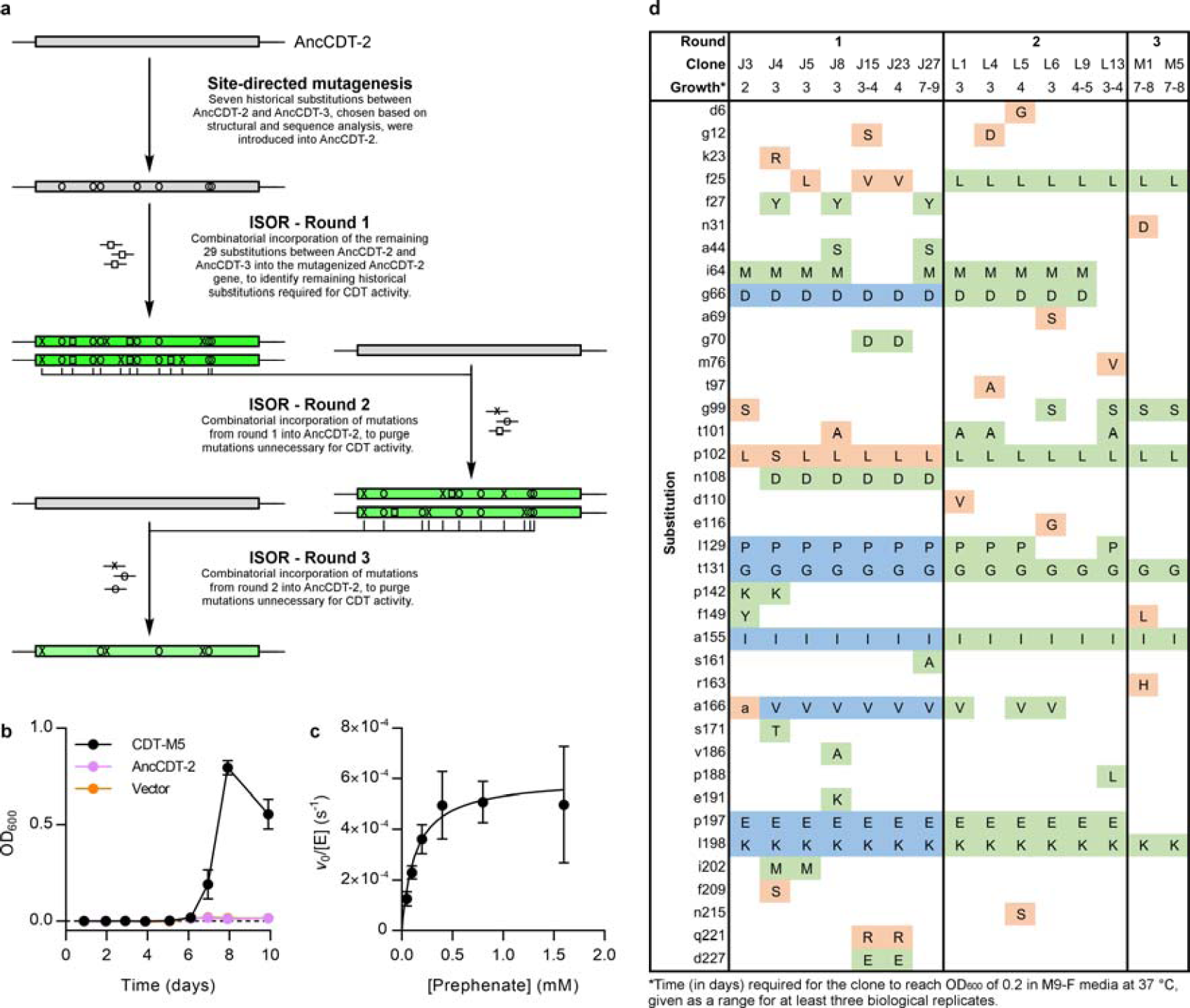
Directed evolution of AncCDT-2. **a**, Overview of the strategy used for directed evolution of AncCDT-2. **b**, Complementation of auxotrophic *E. coli kpheA* cells in selective M9-F media by CDT-M5 (mean ± s.e.m., three biological replicates). AncCDT-2 and empty vector transformants were used as negative controls. **c**, Michaelis-Menten plot for CDT-M5 (mean ± s.d. of 3 – 8 technical replicates). d, Sequences of AncCDT-2 variants with CDT activity, isolated by genetic selection of ISOR libraries. Amino acid substitutions in blue originated from the template gene (*via* site-directed mutagenesis), substitutions in green were encoded in oligonucleotides, and substitutions in orange were acquired randomly.

Although AncCDT-3(P188L) has CDT activity, its second order rate constant (*k*_cat_/*K*_M_) is ∼6000-fold lower than PaCDT, despite their active sites being virtually identical (**Extended Data Figs 3h and 9**). We therefore investigated the role of structural dynamics in the evolutionary process. Upon ligand binding, SBPs undergo domain-scale open-closed conformational changes that are essential for function^13^, exemplified by the unliganded and arginine-bound crystal structures of AncCDT-1 (**Fig. 4a**). The open-closed conformational equilibrium of an SBP controls binding affinity^14^ and the rate of solute transport^13^, suggesting that this equilibrium is subject to evolutionary selection. On the other hand, efficient enzyme catalysis depends on pre-organization of the active site; unproductive conformational sampling has been shown to constrain the catalytic efficiency of recently evolved enzymes^15,16^. The closed conformation of CDT is the catalytically competent conformation; the open-closed conformational change would be necessary only to the extent needed to enable substrate binding and product release from the occluded active site.

**Figure 4.**
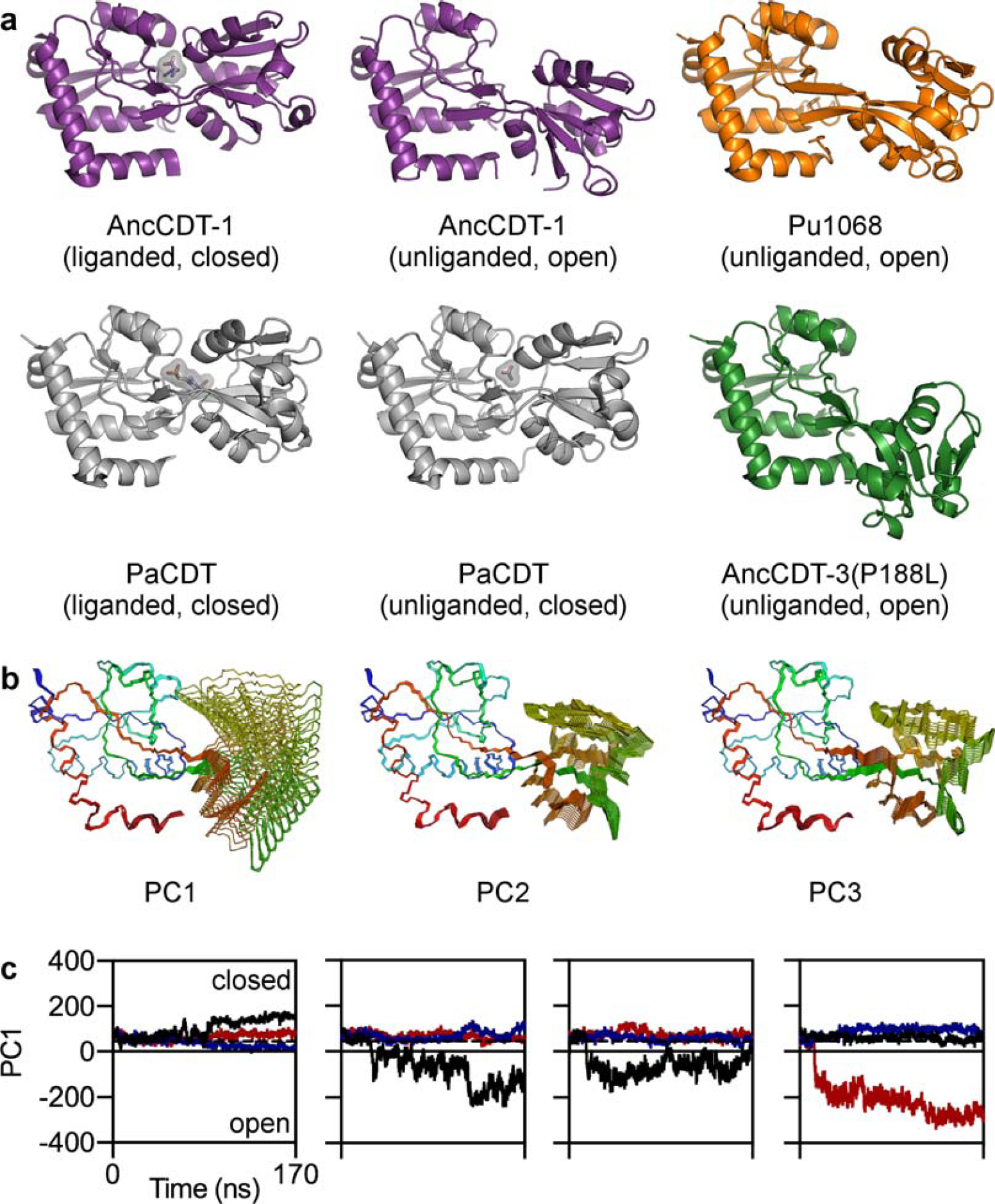
Structural dynamics of CDT. **a**, Open and closed structures of AncCDT-1, Pu1068, PaCDT, and AncCDT-3(P188L). Unusually, PaCDT adopts a closed structure in the absence of ligand. **b**, Principal component analysis of MD simulations of PaCDT. The structures illustrating the physical interpretation of the first three principal components (PCs) were generated by interpolating between structures at the extremities of each principal component axis. These motions represent hinge-bending and hinge-twisting motions typical of AABPs^18,19^. **c**, Open-closed conformational dynamics in 4 × 170 ns simulations of PaCDT, initialized from the unliganded structure (PDB: 5HPQ) using the GROMOS 53a6 force field. Projections of the trajectories of individual PaCDT subunits onto the PC1 axis are shown. Each color represents a subunit of the PaCDT homotrimer. The dotted line represents the crystallographic conformation (PDB: 5HPQ).

The unliganded SBPs AncCDT-1 and Pu1068 and the inefficient ancestral enzyme AncCDT-3(P188L), whose structures were solved in this work, crystallized in an open conformation (**Fig. 4a**). This is consistent with previous studies showing that unliganded AABPs sample closed or semi-closed conformations only transiently^13,17^, and with previously reported crystal structures of unliganded AABPs, of which only 1/14 crystallized in a closed conformation (**Supplementary Table 4**). In contrast, PaCDT crystallized in a closed conformation in the absence of substrate or substrate analogs in multiple, differently packed crystals, suggesting that the closed conformation of the enzyme is unusually stable for this protein fold (**Fig. 4a and Extended Data Fig. 6a**). Molecular dynamics (MD) simulations of the PaCDT trimer initialized from this structure, totaling 680 ns of simulation time, indicated that the open conformation is accessible in PaCDT, although most of the subunits remained closed throughout a 170 ns trajectory (**Fig. 4b–c, Extended Data Fig. 10**). Additional simulations, totaling 550 ns of simulation time, using a different initial structure or a different force field gave similar results (**Extended Data Fig. 10d–e**). The domain-scale conformational fluctuations that did occur in these MD simulations were characteristic of SBPs; principal component analysis showed that hinge-bending and hinge-twisting motions typical of AABPs^18,19^ accounted for >85% of conformational variance (**Fig. 4b**). Indeed, the open structure of AncCDT-3(P188L), which provided experimental evidence for sampling of the open conformation in CDTs, resembled the simulated open conformation of PaCDT (**Extended Data Fig. 10a–c**). Thus, the characteristic domain-scale dynamics of the SBP fold are retained in CDTs and are indeed necessary for substrate/product diffusion from the occluded active site. However, the unusual stability of the closed conformation of PaCDT suggests that the conformational landscape of the enzyme has evolved between AncCDT-3(P188L) and PaCDT to minimize unproductive sampling of the non-catalytic open conformation, contributing to improvements in catalytic efficiency towards the end of the evolutionary trajectory.

**Extended Data Figure 9.**
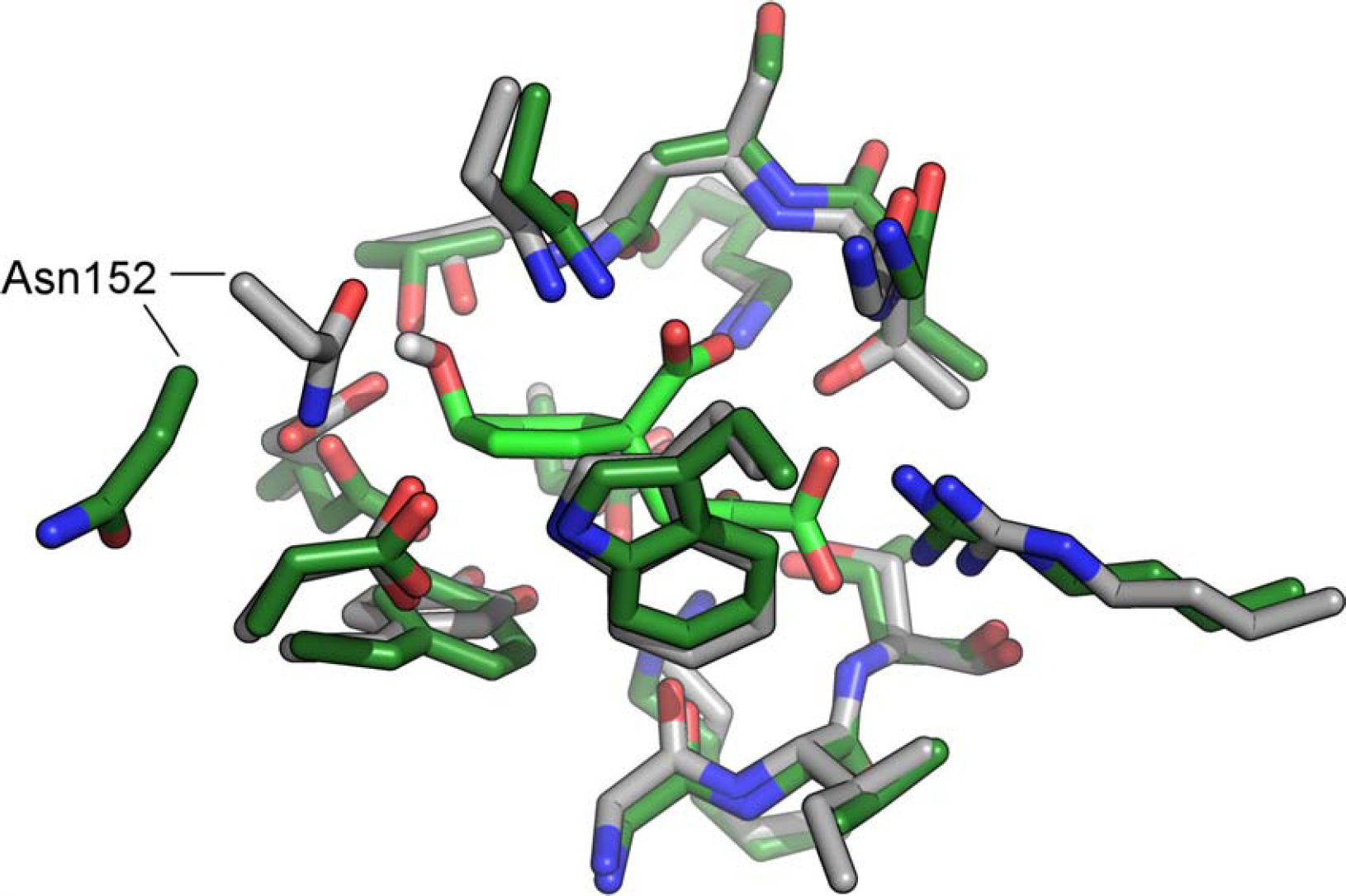
Active site structures of PaCDT and AncCDT-3(P188L). The closed conformation of AncCDT-3(P188L) was modeled by superimposing the two domains of AncCDT-3(P188L) (dark green) separately on the structure of PaCDT (grey, with docked prephenate in light green). Excluding Asn152, which is involved in crystal packing in the AncCDT-3(P188L) structure, the active site structures of the two proteins are virtually identical, despite the difference in global conformation.

**Extended Data Figure 10.**
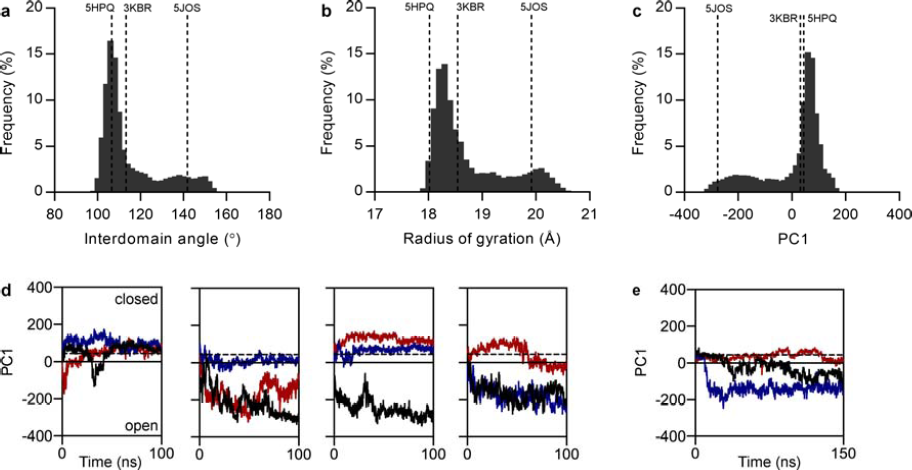
Molecular dynamics simulations of PaCDT. **a-c**, Frequency histograms of the interdomain angle (a), the radius of gyration (b), and the projection onto the first principal component (PC1) (c) for individual PaCDT subunits in the eight simulations using the GROMOS 53a6 force field (1.08 μs simulation time). Each quantity can be used as a descriptor of the conformational change between the open and closed states of the protein. The corresponding values for the crystal structures of PaCDT (PDB: 5HPQ, 3KBR) and AncCDT-3(P188L) (PDB: 5JOS) are also shown. **d-e**, Projections of the trajectories of individual PaCDT subunits onto the PC1 axis. Each color represents a subunit of the PaCDT homotrimer. The dotted line represents the crystallographic conformation (5HPQ). (**d**) 4 × 100 ns simulations initialized from 3KBR using the GROMOS 53a6 force field. (**e**) 1 × 150 ns simulation initialized from 5HPQ using the OPLS3 force field.

Our results suggest that the evolution of highly specialized and efficient CDTs (e.g., PaCDT, *k*_cat_/*K*_M_ ∼ 10^6^ M^-1^ s^-1^) from non-catalytic ancestors occurred in several distinct stages. Incorporation of the desolvated general acid Glu173 into the binding pocket of an ancestral SBP, despite initially being an adaptation for a different function, may have been sufficient for initial, promiscuous CDT activity. Indeed, the intrinsic reactivity of desolvated acidic and basic residues has been exploited similarly in enzymes that have evolved recently in response to anthropogenic substrates^20^ and in enzymes engineered *via* single substitutions in non-catalytic proteins^21^. Following the introduction of a reactive general acid, optimization of enzyme-substrate complementarity and the introduction of hydrogen-bonding networks to position the catalytic residue precisely and stabilize the departing carboxylate group of the substrate appear to have occurred. Further improvements in catalytic efficiency could have been gained by second-and third-shell substitutions that refine the structure of the active site and optimize conformational sampling to favor catalytically relevant conformations. Similar mutational patterns have been documented in directed evolution experiments^15,22^. Additionally, adaptation of protein dynamics has been shown to occur analogously in the evolution of a binding protein from an enzyme, in which case the catalytically relevant conformation was *disfavored* by the function-switching mutation^23^.

Although some computationally designed protein structures have been made with atomic-level accuracy^24^, and various strategies have been developed to introduce catalytic activity into arbitrary protein scaffolds^21,25,26^, replicating the catalytic proficiency of natural enzymes using computational design remains a major challenge^27,28^. The evolutionary trajectory of CDT has striking similarities with the optimization of rationally designed enzymes by directed evolution^29^; catalytic activity can be initialized by computationally guided grafting of a reactive catalytic motif (e.g., a desolvated carboxylate) into a protein scaffold that can accommodate the transition state for a given reaction, and directed evolution can be used to introduce additional stabilizing interactions, optimize positioning of catalytic groups, improve enzyme-transition state complementarity, and optimize conformational sampling, frequently *via* remote substitutions^29,30^. Thus, the strategies that have been used to improve catalytic activity in computational design and directed evolution experiments appear to mirror those that drove the emergence of an enzyme from a non-catalytic protein by natural selection.

## Methods

### Materials

pDOTS7 is a derivative of pQE-82L (QIAGEN) modified to enable Golden Gate cloning^31^, and was created by removal of the *SapI* site from pQE-82L and introduction of two reciprocal *Sap*I sites following the His_6_ tag, with the *Sap*I sites separated by a 28 bp stuffer fragment. This vector was obtained from Prof. Harald Janovjak (IST Austria). The Δ*pheA* strain of *E. coli* K-12 from the Keio collection^32^ (strain JW2580-1) was obtained from the Coli Genetic Stock Center (Yale University, CT).

### Phylogenetic analysis and ancestral sequence reconstruction

The protein sequences of 113 homologs of Ws0279 and PaCDT were collected from the NCBI reference sequence database using the BLAST server. The sequences were aligned in MUSCLE^33^. The alignment was edited to remove N-terminal signal peptides and large insertions, and combined with a subset of a previous alignment of representative AABP sequences^34^ by profile-profile alignment in MUSCLE, which yielded an outgroup of 271 AABP sequences. Phylogenetic trees were inferred using the maximum-likelihood (ML) method implemented in PhyML^35^. Evaluation of BIONJ trees reconstructed using different amino acid substitution models, using the Akaike information criterion as implemented in ProtTest^36^, supported the use of the WAG substitution matrix with gamma-distributed rate heterogeneity, a fixed proportion of invariant sites, and equilibrium amino acid frequencies estimated from the data (WAG+I+Г+F model). Phylogenies were reconstructed in PhyML by optimization of an initial BIONJ tree using the nearest-neighbor interchange and subtree pruning and regrafting algorithms. Robustness of the resulting tree topology to the substitution model was assessed by repeating the analysis using the LG and JTT substitution matrices (LG/JTT+I+Г+F models), and convergence to the ML tree was checked by repeating the analyses with ten randomized initial trees. Although the resulting trees had essentially identical topologies, the tree inferred using the LG+I+Г+F model had the highest likelihood and was therefore taken as the ML tree. Ancestral protein sequences were reconstructed using the empirical Bayes method implemented in PAML^37^. The ancestral sequences AncCDT-1 to 5 were reconstructed using the LG substitution matrix together with the ML tree inferred using the LG+I+Г+F model, and the ancestral sequences AncCDT-1W to 5W were reconstructed using the WAG substitution matrix together with the tree inferred using the WAG+I+Г+F model (Extended Data Fig. 2).

### Cloning and mutagenesis

Codon-optimized synthetic genes encoding the ancestral proteins, Ws0279 (UniProt: Q7MAG0; residues 24–258), Pu1068 (UniProt: Q4FLR5; residues 19–255), Ea1174 (UniProt: K0ABP5; residues 31–268), and PaCDT (UniProt: Q01269; residues 26–268) were cloned into the pDOTS7 vector using the Golden Gate method^31^. Site-directed mutagenesis was achieved using Gibson assembly^38^: gene fragments with ∼30 bp overlap were synthesized by PCR using complementary primers encoding the desired mutation and assembled together with the linearized pDOTS7 vector using Gibson assembly. Successful cloning and mutagenesis was confirmed by Sanger sequencing of the vector insert.

### Protein expression and purification

Proteins were generally expressed in *E. coli* (BL21)DE3 cells, except for enzyme assays, in which case they were expressed in Δ*pheA* cells to exclude the possibility of contamination with endogenous prephenate dehydratase. Cells were typically grown in Luria-Bertani (LB) or Terrific Broth (TB) media at 37 °C to OD_600_ 0.8, induced with 0.5 mM β-D-1-isopropylthiogalactopyranoside and incubated for a further 20 h at 37 °C. Cells were pelleted and stored at −80 °C prior to protein purification. For most applications, proteins were purified under native conditions by nickel-nitrilotriacetic acid (Ni-NTA) affinity chromatography and size-exclusion chromatography (SEC). Cells were thawed, resuspended in equilibration buffer (50 mM NaH2PO4, 500 mM NaCl, 20 mM imidazole, pH 7.4), lysed by sonication, and fractionated by ultracentrifugation (24,200×*g*; 1 hr, 4 °C). The supernatant was filtered through a 0.45 μm filter and loaded onto a 5 mL HisTrap HP column (GE Healthcare) equilibrated with equilibration buffer. The column was washed with 50 mL equilibration buffer and 25 mL wash buffer (50 mM NaH_2_PO_4_, 500 mM NaCl, 44 mM imidazole, pH 7.4), and the target protein was eluted in 25 mL elution buffer (50 mM Na_2_HPO_4_, 500 mM NaCl, 500 mM imidazole, pH 7.4). For ITC experiments, proteins were subjected to on-column refolding during the affinity chromatography step to remove endogenously bound ligands, as described previously^34^. Proteins were concentrated using a centrifuge filter (Amicon Ultra-15 filter unit with 10 kDa cut-off) and purified by SEC on a HiLoad 26/600 Superdex 200 column (GE Healthcare), typically eluting in SEC buffer (20 mM Na_2_HPO4, 150 mM NaCl, pH 7.4). Protein purity was confirmed by SDS-PAGE, and protein concentrations were measured spectrophotometrically using molar absorption coefficients calculated in ProtParam (http://expasy.org/tools/protparam.html).

### Analytical size-exclusion chromatography

The size-exclusion column (HiLoad 26/600 Superdex 200, GE Healthcare) was calibrated using a set of standard proteins (Gel Filtration HMW Calibration Kit, GE Healthcare) in SEC buffer. The partition coefficient (K_av_) of each protein was calculated using the equation *K*_av_ = (*v*_e_ – *v*_o_)/(*v*_c_ – *v*_o_), where *v_e_* is the elution volume, *v*_o_ is the column void volume, and *v*_c_ is the geometric column volume, and used to construct a calibration curve of *K*_av_ versus log(molecular mass).

### Differential scanning fluorimetry

Differential scanning fluorimetry (DSF) experiments to test Ws0279, AncCDT-1, Pu1068, and AncCDT-2 for binding of amino acids and other metabolites were performed using a ViiA 7 (Thermo Scientific) or 7900HT Fast (Applied Biosystems) real-time PCR instrument. Reaction mixtures contained 5 μM protein in DSF buffer (50 mM Na_2_HPO_4_, 150 mM NaCl, pH 7.6), 5×; SYPRO orange dye (Sigma-Aldrich) and ligand (1 mM or 10 mM for amino acids, ≥10 mM for other metabolites) in a total volume of 20 μL, and were dispensed onto a 384-well PCR plate, at least in triplicate. At least eight replicates of ligand-free control were also included on each plate. Fluorescence intensities were monitored continuously as the samples were heated from 20 °C to 99 °C at a rate of 0.05 °C/s, with excitation at 580 nm and emission measured at 623 nm. Melting temperatures (*T*_M_) were determined by fitting the data to a Boltzmann function, *F* = *AT* + B + (C*T* + D)/(1+exp((*T*_M_ – *T*)/E)), where *F* is fluorescence and *T* is temperature. The parameters A and C, accounting for the slopes of the pre-and post-transition baselines, were fixed at zero if possible.

Pu1068, AncCDT-1, and AncCDT-2 were also screened against a subset of Biolog Phenotype Microarray (PM) plates (Biolog, Hayward, CA, USA), as described previously^39^. Libraries of biologically relevant potential ligands were generated by dissolving each compound in 50 μL water, resulting in concentrations of approximately 10–20 mM in the assay (the exact concentrations vary from well to well, and are not released by the manufacturer). Plates PM1–PM5 contain single concentrations of each compound, while plate PM9 contains a series of concentrations of each compound. Fluorescence intensities were measured on a Lightcycler 480 real-time PCR instrument (Roche Diagnostics). Initial hits were further tested using known concentrations (0–600 mM) of each potential ligand to confirm binding. An additional in-house screen consisted of a subset of the Solubility and Stability Screen (Hampton Research), which was tested by the CSIRO Collaborative Crystallisation Centre (http://www.csiro.au/C3), Melbourne, Australia. For this screen, the reaction mixtures contained 0.3 μg Pu1068, 3.75× SYPRO orange and 5 μL ligand in a total volume of 20 μL, in a 96-well plate format; each ligand was tested at three concentrations and three replicates of a ligand-free control were also included. Fluorescence intensities were measured on a BioRad CFX384 real-time PCR instrument with excitation at 490 nm and emission at 570 nm. The temperature was ramped from 20 °C to 100 °C at a rate of 0.05 °C/s, and the fluorescence intensity was measured at 0.5 °C intervals. Melting temperatures were taken as the temperature at the minimum of the first derivative of the melt curve, which was determined by fitting the data to a quadratic function in the vicinity of the melting temperature using GraphPad Prism 7 software.

### Isothermal titration calorimetry

ITC experiments were performed using a Nano-ITC low-volume calorimeter (TA Instruments); details of instrument calibration have been described previously^34^. ITC experiments were performed at 25 °C with stirring at 200 rpm. Protein and ligand solutions were prepared in matched SEC buffer and degassed before use. Amino acid solutions were prepared volumetrically from commercial samples (Sigma-Aldrich, Alfa Aesar) with stated purity ≥98%. Ancestral proteins were tested for binding of proteinogenic amino acids *via* screening experiments in which 45 μL of 0.844 mM ligand was injected continuously into 164 μL of 50 μM protein over 300 s. In some cases, ligands were tested in mixtures of structurally related amino acids. For quantitative titrations, 100 μM protein was generally titrated with 1 × 1 μL, then 28 × 1.6 μL injections of 0.69 mM ligand at 300 s intervals. The background heat was estimated as the average heat associated with each injection in a control titration of ligand into buffer, and subtracted from each protein-ligand titration. Association constants (*K*_a_) were determined by fitting the integrated heat data to the independent binding sites model in NanoAnalyze software (TA Instruments).

### Genetic complementation

*E. coli* strain JW2580-1 (Δ*pheA*) cells were transformed with the appropriate plasmid by electroporation, plated on LB agar supplemented with 100 mg/L ampicillin (LBA agar), and incubated at 37 °C overnight. Single colonies were used to inoculate 20 mL M9 minimal media supplemented with L-tyrosine, ampicillin and IPTG (M9–F; per L: 6 g Na2HPO4, 3 g KH2PO4, 0.5 g NaCl, 1 g NH_4_Cl, 20 mL 20% (w/v) glucose, 2 mL 1 M MgCl_2_, 0.1 mL 1 M CaCl_2_, 2 mL 2.5 mg/mL L-tyrosine, 1 mL 100 mg/mL ampicillin, 0.2 mL 1 M IPTG). The cultures were incubated at 37 °C with shaking at 180 rpm, and OD_600_ was measured periodically. We confirmed that the observed differences in growth rates could not be explained by differences in protein expression by culturing each clone in M9–F media supplemented with 20 μg/mL L-phenylalanine (M9+F media) and assessing protein expression by SDS-PAGE of the soluble fraction of the crude cell lysate from each culture.

### Preparation of sodium prephenate

Sodium prephenate was prepared from barium chorismate (Sigma, 60 – 80% purity). Barium chorismate (40 mM in H2O) was mixed with an equimolar amount of 1 M Na_2_SO4. An equal volume of 100 mM N2HPO4 (pH 8.0) was added to the mixture, and the BaSO_4_ precipitate was removed by centrifugation. Sodium prephenate was obtained by heating the resulting sodium chorismate solution at 70 °C for 1 hr^40^. Aliquots were stored at −80 °C. The concentration of prephenate was measured by quantitative conversion of prephenate to phenylpyruvate under acidic conditions (0.5 M HCl, 15 min, 25 °C) and spectrophotometric determination of phenylpyruvate concentration, as described previously^41^.

### Prephenate dehydratase assay

Prephenate dehydratase activity was determined by spectrophotometric measurement of phenylpyruvate formation, as described previously^41^. Protein solutions were prepared in 20 mM Na_2_HPO_4_, 150 mM NaCl (pH 7.4), and prephenate solutions were prepared in 50 mM Na2HPO4 (pH 8.0). After equilibration at room temperature for 5 min, the reaction was initiated by mixing equal volumes of protein and substrate solutions. Aliquots (50 μL or 100 μL) were regularly removed from the reaction mixture and quenched by addition of an equal volume of 2 M NaOH. Absorbance at 320 nm was measured using an Epoch Microplate Spectrophotometer (BioTek), and phenylpyruvate concentrations were determined assuming a molar extinction coefficient of 17,500 M^−1^ cm^−1^. Reaction times and enzyme concentrations were adjusted to ensure <20% conversion of prephenate to phenylpyruvate. The rate of non-enzymatic turnover was subtracted from the observed rate of enzyme-catalyzed turnover.

### Circular dichroism spectroscopy

Circular dichroism (CD) experiments were performed using a Chirascan spectropolarimeter (Applied Photophysics) with a 1-mm path length quartz cuvette. Proteins were diluted to 0.3 mg/mL in water (for recording CD spectra) or SEC buffer (for thermal denaturation experiments) and degassed prior to measurements. CD spectra were recorded at 20 °C between 190 nm and 260 nm, with a bandwidth of 0.5 nm and a scan rate of 3 s per point, with adaptive sampling. For thermal denaturation experiments, CD was monitored at 222 nm over a temperature range of 20 °C to 90 °C, heating at 1 °C min^-1^. *T*_M_ values were determined by fitting the data to a two-state model:

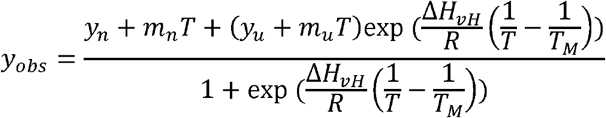

where *y*_obs_ is ellipticity at 222 nm, *y*_n_, *m*_n_, *y*_u_, and *m*_u_ describe the pre-transition and post transition baselines, *T* is temperature, *R* is the gas constant, and ∆H_vH_ is the apparent van’t Hoff enthalpy of unfolding.

### Crystallization and structure determination

Crystal structures of AncCDT-1 (complexed with L-arginine), Pu1068 (unliganded), AncCDT-3(P188L), and PaCDT were solved and refined at resolutions between 1.6 Å and 2.6 Å. An additional low-resolution structure of PaCDT (3.1 Å) shows an alternate crystal packing arrangement, and a low-resolution structure of unliganded AncCDT-1 (3.4 Å) illustrates the domain-scale conformational change resulting from ligand binding. Pu1068 was also co-crystallized with NDSB-221 ((3-(1-methylpiperidinium-1-yl)propane-1-sulfonate); this low-affinity ligand was identified by DSF and confirmed by fluorescence spectroscopy to bind with a K_d_ of 0.53 mM (**Extended Data Fig. 5**).

AncCDT-1, AncCDT-3(P188L), Pu1068, and PaCDT were crystallized using the vapor diffusion method at 18 °C. Crystals were cryoprotected and flash frozen in a nitrogen stream at 100 K. Diffraction data were collected at 100 K on the MX1 or MX2 beamline of the Australian Synchrotron^42^. The data were indexed and integrated in iMOSFLM^43^ or XDS^44^, and scaled in Aimless^45^. Structures were solved by molecular replacement in Phaser^46^ and refined by real space refinement in Coot^47^ and reciprocal space refinement in REFMAC5^48^ and/or PHENIX^49^. Full details of crystallization and structure determination for each protein are given in **Supplementary Tables 5-8**. Data collection and refinement statistics are given in **Supplementary Tables 9-12**.

### Computational docking

The apo-PaCDT structure (PDB: 5HPQ) was prepared for computational docking in Maestro (Schrödinger). Missing side chains were rebuilt. Glu173 was protonated, and other residues were assigned the appropriate protonation state at pH 7.0. Asn, Gln, and His side-chains were flipped, and Ser, Thr, Tyr, and water hydroxyl groups were reoriented to optimize hydrogen bonding networks. The structure was energy-minimized under the OPLS3 force field, with heavy atoms restrained within 0.3 Å of their initial position. Water and acetate molecules were removed from the structure after energy minimization. The structures of the PaCDT/prephenate and PaCDT/L-arogenate complexes were modeled by computational docking in Glide (Schrödinger) using the standard precision mode with default parameters for docking and scoring. The resulting complexes were energy minimized using the OPLS3 force field. In their respective highest scoring poses, L-arogenate and prephenate adopted the expected orientation, with the α-amino acid and α-keto acid moieties binding at the conserved structural motif that recognizes the same functional groups in AABPs.

### Staggered extension process

AncCDT-3 and AncCDT-3W were recombined using the staggered extension process (StEP) following a literature protocol^50^. The StEP reaction mixture contained 5 μL 10× *Taq* buffer, 1.5 mM MgCl_2_, 0.2 mM each dNTP, 75 fmol each template plasmid, 30 pmol each primer, and 2.5 U *Taq* polymerase (New England Biolabs) in a total volume of 50 μL. The primers used in the reaction were the 5´ flanking primer P7XF and the 3´ flanking primer P7XR (**Supplementary Table 13**), which amplify ∼100 bp on either side of the *SapI* site of the pDOTS7 vector. The thermocycling program consisted of 80 cycles of (i) a denaturation step for 30 s at 95 °C; and (ii) an annealing/extension step for 5 s at 52 °C. 2 μL of the resulting PCR product was incubated with 10 U *DpnI* (Thermo Scientific) in a reaction volume of 10 μL at 37 °C for 1 hr to digest the parental plasmid DNA. 5 μL of the DpnI-digested StEP product was then amplified in a nested PCR reaction using *Taq* polymerase, in a total volume of 100 μL. The primers used for the nested PCR reaction, P7NF and P7NR (**Supplementary Table 13**), target the *EcoRI* site on the 5´ strand and the *HindIII* site on the 3´ strand of the pDOTS7 vector, respectively. The nested PCR product was run on a 1% agarose gel and purified by gel extraction.

### Incorporation of synthetic oligonucleotides *via* gene reassembly

Incorporation of synthetic oligonucleotides *via* gene reassembly (ISOR) was achieved following literature protocols^51,52^. The template gene was amplified by PCR using Phusion Hot Start II Polymerase (Thermo Scientific) using the primers P7XF and P7XR (Supplementary Table 13). The purified PCR product was digested with DNAse I (New England Biolabs) in a reaction mixture containing 100 mM TRIS pH 7.5, 10 mM MnCl_2_, 4 μg PCR product and 0.3 U DNAse I in a total volume of 40 μL. The reaction mixture was incubated at 37 °C for 1 – 2 min and quenched by the addition of 20 μL 0.1 M EDTA pH 8.0 pre-incubated at 80 °C, followed by heat inactivation at 80 °C for 15 min. The digested PCR product was run on a 2% agarose gel, and fragments 50 – 250 bp in size were excised from the gel and purified using the Wizard SV Gel and PCR Clean-Up System (Promega). The fragments were reassembled using *Taq* polymerase: each reaction contained 40 ng gene fragments, 2 μL 10×; buffer, 0.2 mM dNTPs, 1.25 U *Taq* polymerase and varied concentrations of equimolar mutagenic oligonucleotides (5 – 800 nM total concentration) in a volume of 20 μL (see **Supplementary Table 13** for a list of oligonucleotides included in each round). The thermocycling protocol consisted of (i) an initial denaturation step at 95 °C for 2 min; (ii) 40 cycles of a denaturation step at 95 °C for 30 s, then 13 hybridization steps from 65 °C to 41 °C in 2 °C steps, each for 90 s (total 13.5 min), then an extension step at 72 °C for 1 min; and (iii) a final extension step at 72 °C for 7 min. 0.5 μL of the unpurified assembly reaction mixture was amplified in a 50 μL nested PCR reaction using *Taq* polymerase and the primers P7NF and P7NR (**Supplementary Table 13**). The nested PCR product was run on a 1% agarose gel and purified by gel extraction.

### Library creation and selection

Purified PCR products (0.5 μg) from StEP or ISOR reactions were digested with 2.5 μL each of *HindIII* FD and *EcoRI* FD (Thermo Scientific) in a 50 μL reaction at 37 °C for 30 min. The reaction mixture was purified immediately using a PCR purification kit. The pDOTS7 vector containing the AncCDT-2 insert (2.5 μg) was digested using 2.5 μL each of *HindIII* FD, *EcoRI* FD, and *Psf*I FD (which cuts within the AncCDT-2 insert) in a 50 μL reaction at 37 °C for 30 min. The digested vector was purified immediately using a PCR purification kit, then run on a 1% agarose gel and purified by gel extraction. Ligation reaction mixtures contained 100 ng pDOTS7 vector, a 3-fold molar excess of insert, 2 μL 10× T4 DNA ligase buffer, and 5 U T4 DNA ligase (Thermo Scientific) in a volume of 20 μL, and were incubated at room temperature for 1 hr. Following purification of the ligation reaction mixture using a PCR purification kit, electrocompetent *E. coli* strain JW2580-1 (Δ*pheA)* cells were transformed with 1 μL ligation product by electroporation and plated on LBA agar. Following overnight incubation of the plates at 37 °C, colonies were scraped into LB media, then resuspended in 20 mL fresh LBA media. 100 μL of the resulting cell suspension was used to inoculate 20 mL fresh LBA media, which was then incubated at 37 °C until the OD600 reached ∼0.5. A 1 mL aliquot of the culture was washed twice with 1 mL M9 salts (6 g/L Na_2_HPO_4_, 3 g/L KH_2_PO_4_, 1 g/L NH_4_Cl, 0.5 g/L NaCl), and resuspended in 1 mL M9 salts. Serial dilutions of the cell suspension were made in M9 salts, plated on M9–F agar, and incubated at 37 °C. The resulting colonies were streaked onto LBA agar, and their plasmid DNA was amplified by PCR using the sequencing primers P7SF and P7SR (**Supplementary Table 13**). The resulting PCR products were sequenced by GENEWIZ (South Plainfield, N.J., U.S.A.) or the Biomolecular Resource Facility at ANU. Single colonies from the streaked LBA plates were used to confirm growth of the clone in liquid M9–F media, as described above, and to inoculate LBA cultures, from which plasmid DNA was extracted.

### Molecular dynamics simulations

MD simulations were initialized from the HEPES-bound and unliganded PaCDT structures (PDB: 3KBR, 5HPQ). The structure of PaCDT trimer was generated from the monomer structure by application of the crystallographic three-fold rotation operation. Small molecules were removed from the structures, and missing side-chains and a missing residue (Gln190) in the HEPES-bound structure were modelled in MODELLER^53^. N-terminal acetyl caps and C-terminal amide caps were added using MODELLER and Coot^47^. MD simulations were performed using GROMACS version 4.5.5 (ref. ^54^) for the HEPES-bound structure and GROMACS version 4.6.5 for the unliganded structure, using the GROMOS 53a6 force field^55^ in both cases. The protein was solvated in a rhombic dodecahedron with SPC water molecules, such that the minimal distance of the protein to the periodic boundary was 15 Å, and 15 Na^+^ ions were added to neutralize the system. Energy minimization was achieved using the steepest descent algorithm. A 100 ps isothermal (NVT) MD simulation with position restraints on the protein was used to equilibrate the system at 300 K. For production MD simulations of the NPT ensemble, the temperature was maintained at 300 K using Berendsen’s thermostat (τ_T_ = 0.1 ps), and the pressure was maintained at 1 bar using Berendsen’s barostat (τ_p_ = 0.5 ps, compressibility = 4.5 × 10^-5^ bar^-1^). All protein bonds were constrained with the LINCS algorithm; water molecules were constrained using the SETTLE algorithm; the time step for numerical integration was 2 fs; the cut-offs for short-range electrostatics and van der Waals forces were 9 Å and 14 Å, respectively; the Particle-Mesh Ewald method was used to evaluate long-range electrostatics; neighbor lists were updated every 10 steps. Following a 1 ns equilibration phase, which was not considered in the analysis, the four simulations of the HEPES-bound structure were continued for 100 ns, and the four simulations of the unliganded structure were continued for 170 ns.

An additional 150 ns simulation was performed in Desmond version 4.8 (Schrödinger 2016-4) (ref. ^56–58^) using the OPLS3 force field^59^. Simulations were initiated from the same starting structure used in the 5HPQ GROMOS simulations, except that Desmond was used to add the N-terminal acetyl caps and C-terminal amide caps, and for energy minimization of the protein structure. The protein was solvated in an orthorhombic box (15 Å periodic boundary) with TIP3P water molecules. Na^+^ ions were added to neutralize the system. Energy minimization was achieved using the steepest descent algorithm (2000 iterations and a convergence threshold of 1 kcal/mol/Å). The system was relaxed using the default Desmond relaxation procedure at 300 K. For production MD simulations of the NPT ensemble, the temperature was maintained at 300 K using a Nosé-Hoover thermostat (τ_T_ = 1.0), and the pressure was maintained at 1.01 bar (τ_p_ = 2.0) using a Martyna-Tobias-Klein barostat. Otherwise, default Desmond options were used. Following equilibration (160 ps), the simulation was run for 150 ns.

### Structural analysis

Residues in extant CDT homologs (Ws0279, Pu1068, Ea1174, PaCDT) are numbered according to the equivalent position in the ancestral proteins. Bio3D^60^ was used for root-mean-square deviation, radius of gyration, and interdomain angle calculations, and principal component analysis. These analyses were performed on the 3KBR and 5HPQ-GROMOS simulations using protein backbone atoms (N, C, Cα) of individual protein subunits at 0.1 ns intervals. The 5HPQ-OPLS simulations were analyzed separately and projected onto the principal components derived from the 3KBR and 5HPQ-GROMOS simulations. The interdomain angle was calculated as the angle between the centers of mass of three groups of backbone atoms: the large domain (residues 2–97 and 196–234), the hinge region (residues 96-98 and 196–198) and the small domain (residues 98–195). Hinge axes for rigid-body domain displacements were determined using DynDom^61^ (**Extended Data Fig. 6d**). PROPKA^62^ was used for pKa prediction.

### Intrinsic tryptophan fluorescence spectroscopy

Intrinsic tryptophan fluorescence spectra were recorded using a Cary Eclipse fluorimeter. Pu1068 was prepared at a concentration of 5 μM in DSF buffer. The excitation wavelength was 280 nm, and emission was measured between 300 nm and 400 nm. Following addition of each aliquot of NDSB-221, the sample was incubated at ambient temperature for 1 min before the fluorescence spectrum was recorded. The *K*_d_ for the Pu1068/NDSB-221 interaction was calculated by fitting the fluorescence data to a hyperbolic binding curve: *F* = *F*_0_ + (*F*_max_ − *F*_0_) × [L]/(*K*_d_ + [L]), where *F* is fluorescence, *F*_0_ and *F*_max_ are initial and final fluorescence, and [L] is ligand concentration.

## Acknowledments

B.E.C. and J.A.K. were supported by Australian Postgraduate Awards. B.E.C. was also supported by a Rod Rickards PhD Scholarship and an Alan Sargeson scholarship. This research was undertaken with the assistance of resources, services, and staff from the Australian National Computational Infrastructure (NCI) and the Australian Synchrotron, and funding from the Australian Research Council Discovery Project scheme (C.J.J.). We thank A. Saeed, P. Yates, L. Tan and S. L. Warring for additional technical contributions.

## Author contributions

B.E.C. and C.J.J. conceived the study; B.E.C. performed computational analysis; J.A.K., B.E.C., and M.L.G. performed experimental characterization of proteins; B.E.C., J.A.K., P.D.C. and C.J.J. solved the crystal structures; N. T. and C. J. J. supervised students; B.E.C., J.A.K., and C.J.J. wrote the paper. All authors contributed to experimental design, editing the paper, and interpretation of results.

## Author information

Crystal structures have been deposited in the Protein Data Bank under accession codes 5HPQ (PaCDT, space group H3), 5JOT (PaCDT, space group P4_3_22), 5HMT (Pu1068, unliganded), 5KKW (Pu1068, NDSB-221 complex), 5TUJ (AncCDT-1, unliganded), 5T0W (AncCDT-1, L-arginine complex), and 5JOS (AncCDT-3(P188L)). The authors declare no competing financial interests. Correspondence and requests for materials should be addressed to C.J.J.

